# Bioabsorbable Magnesium Metal Scaffolds Improve Dermal Wound Healing and Tissue Regeneration

**DOI:** 10.64898/2026.03.03.709352

**Authors:** Mary Elizabeth Guerra, Nabila N. Anika, Ajeet Nagi, Tracy M. Hopkins, Xiaoxian An, Ling Yu, Pei Liu, Chihtong Lee, Sundeep G. Keswani, Raudel Avila, Sarah K. Pixley, Swathi Balaji

**Author notes:** Corresponding authors: Sarah Pixley, PhD, Department of Pharmacology, Physiology, and Neurobiology University of Cincinnati College of Medicine, 231 Albert Sabin Way, MSB 4206A Cincinnati, OH 45267-0576; Swathi Balaji, PhD Feigin Center, C.450.05 1102 Bates Avenue, Houston, TX 77030, Texas Children’s Hospital and Baylor College of Medicine. Equally contributing authors (Mary Elizabeth Guerra, Nabila N. Anika).

## Abstract

**Objective:** Evaluate the effects of bioabsorbable magnesium wires on dermal wound healing and tissue regeneration in a murine full-thickness wound model.

**Approach:** 6 mm diameter stented dorsal skin wounds were created in C57BL/6J mice and treated with implanted WE43B magnesium alloy wires or PBS control. Wound healing was evaluated on days 7 and 28 by histology, immunohistochemistry, and micro-CT. Finite element analysis modeled mechanical strain distribution due to wire degradation during healing.

**Results:** At day 7, magnesium wire-treated wounds showed 100% improved granulation tissue formation, reduced inflammation (37% fewer CD45+ leukocytes and 37% fewer F4/80+ macrophages), increased neovascularization (91% more CD31+ lumens), and 74% more nerve bundles. Improved wound closure (mean difference -1.48 mm) did not reach statistical significance (d = 1.06). By day 28, magnesium-treated wounds showed improved collagen organization and normalized epidermal thickness. The increase in dermal appendages (247%), and vascular density (41%) did not reach statistical significance. Micro-CT confirmed progressive wire degradation. Modeling revealed that degrading wires dynamically altered strain gradients in healing tissue, thereby modulating the spatial mechanical cues that govern fibroblast migration and extracellular matrix (ECM) remodeling.

**Innovation:** Magnesium is an essential trace element involved in cellular processes critical to wound repair, including angiogenesis, nerve growth, inflammation modulation, and ECM remodeling. Previous magnesium delivery systems incorporated polymers or other confounding materials that degrade rapidly. We directly applied bioabsorbable pure magnesium metal to provide sustained ion release and favorable mechanical properties to support regenerative healing.

**Conclusion:** Bioabsorbable magnesium wires support regenerative wound healing by reducing inflammation, enhancing neovascularization, and promoting favorable ECM remodeling without adverse inflammatory reactions. These findings provide a strong rationale to harness magnesium metal use in wound healing applications.

## INTRODUCTION

Dermal wound healing is a complex process encompassing hemostasis, inflammation, proliferation, and remodeling.^1^ Given clinical observations of impaired wound healing in patients with trace element deficiencies, the involvement of metal elements in the wound healing phases has been well studied.^2^ Magnesium is a cofactor for over 600 enzymes involved in cellular functions essential to repair, including angiogenesis, nerve regeneration, and extracellular matrix (ECM) remodeling.^2–12^ Its degradation also generates hydrogen gas and hydroxide radicals, creating an antimicrobial environment while exerting antioxidant effects.^13,14^

Previous magnesium-based wound therapies have primarily incorporated polymers or composite scaffolds, which introduce additional materials that can confound outcomes and degrade rapidly.^8,15–20^ Therefore, we aimed to optimize the delivery of pure magnesium by using magnesium metal or alloys with minimal impurities to ensure relatively slower release continuously across the wound surface. We previously characterized bioabsorbable magnesium alloy (WE43B) wires in subcutaneous implantation and pure magnesium wires in nerve repair; both deliver sustained magnesium release over weeks, maintain flexibility in tissue, and have demonstrated biocompatibility in tissues.^18,21–27^ Here, we evaluated the capacity of the WE43B alloy wires to promote regenerative healing in murine full-thickness dermal wounds and applied finite element analysis (FEA) to quantify how wire degradation alters local strain gradients during healing. We hypothesized that implanted magnesium wires would expedite wound closure by reducing inflammation, enhancing neovascularization, and promoting neuronal growth.

## INNOVATION

Prior magnesium-based pre-clinical wound therapies relied on hydrogels or composite scaffolds, which introduce additional materials that can confound outcomes and degrade rapidly.^8,15–20^ Here, we directly applied bioabsorbable magnesium metal wires that release ions gradually and mechanically integrate with healing tissue. This dual biochemical and biomechanical approach reduced inflammation, enhanced neovascularization, and normalized scar architecture *in vivo*. Finite element modeling further revealed dynamic strain gradients generated by wire degradation, linking magnesium’s mechanical and immunoregulatory roles. These findings introduce degradable magnesium scaffolds as a novel, clinically translatable platform for regenerative wound healing and surgical tissue repair.

## CLINICAL PROBLEM ADDRESSED

Chronic and acute post-surgical wounds represent a major clinical challenge, affecting millions of patients and contributing to billions in annual healthcare costs.^28–31^ Current wound care materials often fail to actively modulate inflammation or support regenerative tissue architecture, leading to delayed healing and excessive scarring. Magnesium is an essential element with known pro-healing and anti-inflammatory properties, yet its clinical use has been limited by rapid degradation in polymer-based systems.^8,15–20^ This study demonstrates that bioabsorbable magnesium metal scaffolds can provide sustained biochemical and mechanical cues to promote regenerative healing, offering a promising translational platform for advanced surgical and chronic wound management.

## MATERIALS AND METHODS

### Magnesium Wire Preparation

WE43B magnesium alloy wires were produced as described previously and supplied by Ft. Wayne Metals Research Products (Ft. Wayne, IN).^22^ Briefly, wires were cold drawn to 127 µm ø to achieve 90% cold work and then heat treated at 250°C. Wires were cut to 5 mm length and sterilized.^22^ WE43 magnesium contains Y, Nd, Zr, and Gd at 0.1 to 4 wt% (total 6.5 wt%); Ce, Mn, and Zn at <0.1 wt% (total 0.08 wt%), and Cu, La, Fe, Pr, and Ni each at <0.006 (total 0.018 wt%), as described by Soltan et al.^32^ Corrosion for WE43 alloys has been measured but corrosion rates vary significantly between conditions (dry, in solution, *in vivo*) and are further varied *in vivo* due to physical stresses like animal movement.^22,32–34^ In our studies, calculation of corrosion rates was not done because the wires were too small to weigh or were impossible to extract from tissues.^22^

### Animal Studies

C57BL/6J (WT) mice were purchased from Jackson Laboratory (Bar Harbor, ME), bred, and maintained under specific pathogen-free conditions with access to food and water *ad libitum* in the Cincinnati Children’s Hospital Medical Center animal facility. Protocols for animal use were approved by the Cincinnati Children’s Hospital Medical Center Institutional Animal Care and Review Committee.LL

### Wound Healing Model

Mice were anesthetized with inhaled 3% isoflurane. Dorsal skin was shaved and prepared with betadine and isopropyl alcohol. Two full-thickness excisional wounds were created, one on each flank on the dorsum of 8-week-old C57BL/6J WT female mice using a 6-mm punch biopsy (Miltex, Plainsboro, NJ). Wounds were splinted as previously described to prevent immediate contraction and allow for healing by secondary intention.^35^ Five magnesium wires were placed on one of the bilateral wounds, running lengthwise in the animal, and PBS was used on the contralateral side as a control. Wounds were covered with a sterile adhesive dressing (Tegaderm^TM^ 3M, St. Paul, MD) to promote healing in a moist, protected environment **(Supplemental Figure S1a).** Animals received standard pain medication and were monitored daily for the first seven days post-surgery for any signs of distress due to wire placement. All stents were removed on day 7. Mice were euthanized and wounds were harvested at days 7 and 28 for analysis. Six mice were included at each time point.

### Histology and Wound Analysis

Wound tissue, including the wound bed, wound edge, and adjacent uninjured skin were harvested from the euthanized mice and fixed in 10% neutral buffered formalin. The wounds were bisected in half and paraffin embedded. Sections were cut to 5 μm thickness and mounted onto positively charged slides and then deparaffinized and rehydrated following standard protocols prior to staining. All measurements were made by personnel blinded to condition groups.

Wound healing response at day 7 and scar tissue morphology at day 28 post-wounding were determined by hematoxylin and eosin (H&E) and Mason’s trichrome staining, respectively. Epithelial gap and granulation tissue at day 7 were quantified using LASx (Leica application suite) or ImageJ (National Institutes of Health, Bethesda, MD). Epithelial gap was measured as the gap between the two encroaching epithelial wound margins of the newly forming epidermis at day 7 **(Supplemental Figure S1b)**. Granulation tissue at day 7 and scar area at day 28 were measured as the area beginning from the surface at one end of the original wound margin, down to the ipsilateral panniculus carnosus, across the wound cleft to the opposite original wound margin, and finally back up to the skin at the initial wound margin, as described previously.^36^

At day 28, epidermal thickness of the repaired wound (height of the epidermis from the basement membrane of the dermal-epidermal junction to the top cornified layer) was measured at three different points within a high-powered field (HPF, 40x objective) and averaged. Dermal appendages including hair follicles and sebaceous glands per HPF and CD31-positive lumens per HPF were counted in day 28 wounds. In the day 28 wounds, measurements of epidermal thickness and dermal appendages were taken from both the scar tissue and from normal skin at least 2 mm away from the previously wounded area.

ECM architecture and collagen alignment of the repaired wounds at day 28 was assessed by Masson’s Trichrome Staining (Leica Biosystems, Buffalo Grove, IL).^37^ Picrosirius red stain (Polysciences, Warrington, PA) was used to display types of collagen and collagen density. Polarized images were taken of the scar area of the dermis and the area was divided into four regions of interest (ROI) to represent the center and two edges. Within each ROI, the ratio of picrosirius-positive pixels to total pixels in the ROI was quantified to determine the amount of collagen in the selected scar area.

To quantify subepithelial nuclear density, five sampling boxes (34.1 μm^2^) were evenly spaced within each 200 μm section. At least one corner of each sampling box was at the basal edge of the epithelium (areas with appendages or out-of-focus regions were omitted). Using the same procedure, counts were made in two randomly selected 200 μm sections of normal skin (both left and right sides of the scar). ImageJ was used to count fibroblast nuclei per box, including nuclei only if they were completely within the box or touching the left or top borders.

The morphometric differences in epithelial gap, granulation tissue area, and scar area were assessed by different observers. Histologic selections within different HPFs of other parameters were randomly selected to be quantified by multiple observers.

### Immunohistochemistry

Neovascularization (CD31), leukocyte and macrophage infiltration (CD45 and F4/80), and neurons (Tuj1 antibody against neuron-specific tubulin) were assessed with immunohistochemistry (IHC) staining. IHC staining was performed on a Dako Auto-Stainer Link 48 (DakoLink version 4.1, edition 3.1.0.987; Agilent, Santa Clara, CA). Primary antibodies against CD31 (PECAM-1) for endothelial cells (D8V9E; 1:50; Cell Signaling Technology, Danvers, MA), CD45 for pan-leukocytes (ab10558; 1:5000; Abcam, Cambridge, MA) and F4/80 for pan-macrophages (ab111101; 1:100; Abcam, Cambridge, MA) were detected by EnVision+System-HRP (DAB) kits (Dako North America, Carpinteria, CA) and hematoxylin counterstaining. Immunofluorescence staining was done for Tuj1 for nerves (GT11710; 1:500; GeneTex, Irvine, CA) with localization with anti-mouse antibody conjugated with Alexa Fluor™ 594 (A11032; 1:200; Life Technologies, Eugene, OR). Histology slides were imaged with Leica DM 2000® with Leica Application Suite X® version 3.0.4.16529 and Keyence. Percentage of positive cells per HPF within four to six HPFs across the granulating wound bed were quantified for CD45, F4/80, and Tuj1. CD31 staining was used to count the number of blood vessel lumens per HPF for vessel quantification. CD31 staining was also used to quantify vessel lumens in 200 μm^2^ areas proximal or distal to the magnesium wires **(Supplemental Figure S1c)**.

### Micro-Computed Tomography (Micro-CT) Imaging

To assess wire degradation, skin with wounds containing magnesium wires were imaged via micro-CT on the days of sacrifice after a short fixation period and removal to PBS. Imaging was done in a Siemens Inveon Multimodality System (San Diego, CA) in the University of Cincinnati Vontz Core Imaging Laboratory. Samples were scanned at half-degree increments for 192°. Images were acquired with high magnification and a pixel matrix binning of two, resulting in an effective voxel size of 17.27 μm, using 80 kVp voltage and 300 µA current, with the exposure time at 2100 ms with 25 ms settle time. All tissues were kept moist in PBS during the one hour scan, with up to three skin samples placed in a single Styrofoam boat, as shown previously.^21^ Image analysis and preparation of figures was done with either the Inveon software or ImageJ (NIH), using either 3D reconstructions or single frames. Using the Inveon software, tissues with similar Hounsfield Unit radiodensities were selected and either measured or cut out into a new file and rendered into 3D images.

### Modeling of Wound Healing Mechanics with Magnesium Wires

To investigate the influence of the implanted magnesium wires on the mechanical response of the wound during healing, the commercial software ABAQUS (ABAQUS Analysis User’s Manual, 2023) was used for modeling. A coupled mechanical and reactive-diffusion model, implemented under quasi-static conditions, simulated the wound’s response to uniaxial strain (ε*_app_* = 10%) and bending deformations representative of dynamic mouse motions. The tissue structure was meshed with second order modified tetrahedral elements (C3D10M), using approximately 650,000 3D elements to ensure convergence and accurate results. Mesh refinement studies confirmed that strain energy results converged beyond a mesh density of ∼500,000 3D elements. The mouse tissue was modeled as a two-layer structure consisting of an epidermis-dermis (ED) skin layer (*t_ED_* = 0.5 mm) and an underlying subcutaneous tissue (ST) layer (*t_ST_* = 1.0 mm), with an initial wound diameter *d_W_* of 6 mm. Both tissue layers were modeled as nearly incompressible Mooney–Rivlin hyper elastic solids, characterized by the material parameters C_10_ = 13.24 kPa, C_01_ = 3.06 kPa, D_1_ = 0.012 kPa^-1^ for the epidermis-dermis layer (*E*_ED_ = 99 kPa, *v*_ED_ = 0.49) and C_10_ = 16.26 kPa, C_01_ = 4.10 kPa, D_1_ = 0.001 kPa^-1^ for the subcutaneous tissue layer (*E*_ST_ = 122 kPa, *v*_ST_ = 0.49).^38,39^ The five equidistant magnesium wires were modeled as a linearly elastic material with a Young’s modulus *E*_Mg_ = 42,000 MPa and a Poisson’s ratio *v*_Mg_ = 0.34. A parametric study was conducted to quantify the effects of wire geometry and positioning on maximum strain in magnesium, by varying the magnesium wire radius (0.53 - 0.84 mm) and inter-wire spacing (0.9 - 1.4 mm). The concentration-based degradation of the magnesium wires within the wound was modeled as a non-linear reduction in structural stiffness, represented by a decrease in the Young’s modulus *E*_Mg_, due to change in concentration originated by resorption with the surrounding tissue and biofluid as

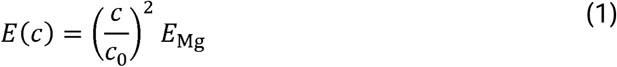

where *c* is the concentration of the magnesium wires over the healing period, and *c*_0_ is the initial concentration of magnesium in the wound.^40^ The concentration over time *c(t)* is modeled as a first-order reaction kinetics following

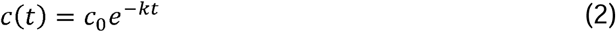

where, *k* = 1.5 s^-1^ is a reaction velocity constant and *t* is the healing time.

### Statistical Analysis

Paired t-tests were used in the comparison of magnesium- and PBS-treated wounds (Prism 10, GraphPad Software, San Diego, CA). A *P* value of < .05 was considered statistically significant. Data are expressed as mean ± standard deviation. Mean differences with corresponding 95% confidence intervals (CI) are also provided. In addition to reporting *P* values, effect sizes were calculated using Cohen’s d (d). Individual data points are shown as dots on quantification figures and represent measurements from independent wounds per animal. Sample size calculations were based on comparisons of PBS and magnesium-treated wounds that yielded N=6 as the total sample size for the two groups (i.e. 3 in each group to identify differences).

An electronic laboratory notebook platform was not used.

## RESULTS

### Magnesium Treatment Improved Wound Closure and Granulation Tissue Formation

All mice tolerated magnesium wire placement well, were active immediately upon post-surgical recovery, groomed themselves, and consumed food and water *ad libitum*. The mouse morbidity score (from zero to two) was not different in mice receiving wires compared to other mice with both dorsal wounds treated with PBS only, as observed on alternate days until day 7 post-wounding. There was no excess inflammatory exudate noted in the magnesium wire-treated wounds compared to the PBS controls and minimal foreign body response (FBR) was observed at day 7 in the magnesium wounds **(Figure 1a-b)**. Wound analysis showed robust connective tissue with minimal collagenous deposition (encapsulation) around the wires. Micro-CT of day 7 wounds showed that the wires were mostly intact and in place within the wounds, although movement of the wires relative to one another was observed and varied from wound to wound **(Figure 1c)**.

**Figure 1.**
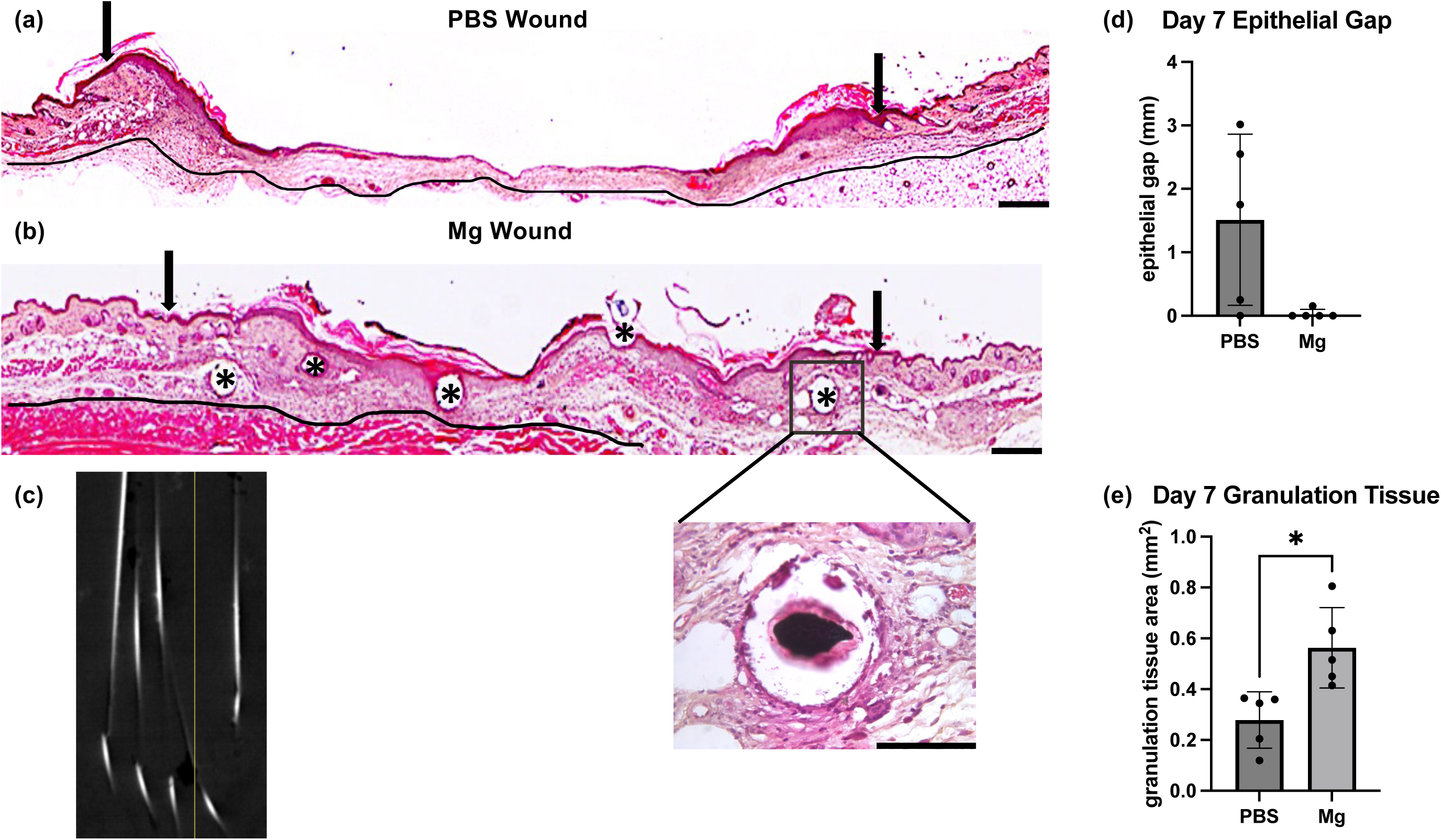
Mg improves wound closure and granulation tissue deposition. (a) H&E staining of PBS wound at day 7 post-wounding. (b) H&E staining of Mg wire-treated wound at day 7 post-wounding. Scale bar = 250 µm. Arrows represent original wound margins. Black lines indicate panniculus carnosus muscle. Asterisks (*) represent locations of magnesium wires, with zoomed in panel (20x) showing minimal FBR around the magnesium wire in the wound. Scale bar = 100 µm. (c) Mg wires in skin explants on micro-CT at day 7. (d) Quantification of epithelial gap at day 7 post-wounding. (e) Quantification of granulation tissue area at day 7 post-wounding. Bar graphs show mean ± SD. Individual dots represent measurements from wounds from different animals. \**P* < .05, \*\**P* < .01, \*\*\**P* < .001.

The effect of magnesium wires on the rate of epithelial gap closure and the amount of granulation tissue in the wounds was then determined. At day 7 post-wounding, magnesium wire treatment resulted in closure of the epithelial gap compared to PBS controls (Mg 0.03 ± 0.07 vs PBS 1.51 ± 1.28 mm; mean difference -1.48 mm; 95% CI -3.21 – 0.25; *P* = .08, d = 1.06) and significantly improved granulation tissue area formation compared to PBS controls (Mg 0.56 ± 0.16 vs PBS 0.28 ± 0.11 mm^2^; mean difference 0.28 mm^2^; 95% CI 0.10 – 0.46; *P* = .01) **(Figure 1d-e)**.

### Magnesium Treatment Reduced Inflammation

To determine the effect of magnesium wire treatment on wound inflammation, we stained day 7 wound cross-sections for CD45, a pan-leukocyte marker, and F4/80, a pan-macrophage marker, which allowed us to assess the level of wound infiltration by leukocytes or macrophages. There was a marked reduction in inflammation in magnesium-treated wounds, as evidenced by significantly fewer CD45-positive cells in magnesium-treated wounds compared to PBS wounds (Mg 21.36 ± 4.31 vs PBS 34.10 ± 5.09 % cells per HPF; mean difference -12.74 % cells per HPF; 95% CI -20.62 – -4.87; *P* = .01) **(Figure 2a-b),** and fewer F4/80-positive cells in magnesium-treated wounds compared to PBS wounds (Mg 31.34 ± 10.12 vs PBS 50.13 ± 7.51 %cells per HPF; mean difference -18.79 % cells per HPF; 95% CI -29.76 – -7.83; *P* < .01) (Figure 2c-d).

**Figure 2.**
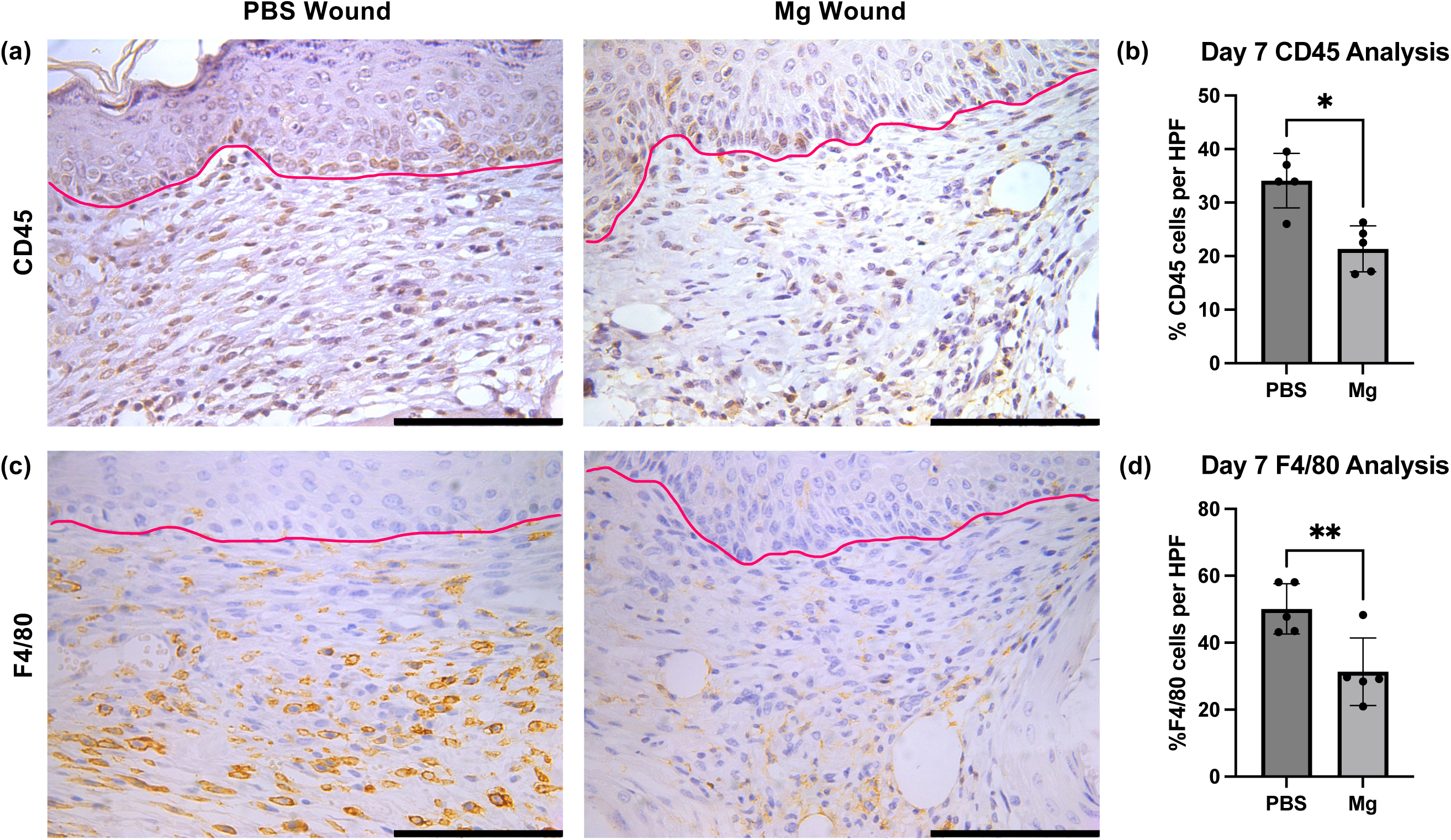
Mg reduces inflammation. CD45 (leukocyte marker) and F4/80 (macrophage marker) IHC staining of Mg and PBS wounds at 7 days post-wounding. (a) CD45 staining representative image of wounds. (b) Quantification of % of CD45 positive cells per HPF (40x) at day 7 post-wounding. (c) F4/80 staining representative image of wounds. (d) Quantification of % of F4/80 positive cells per HPF (40x) at day 7 post-wounding. Scale bar = 100 µm. Pink lines represent dermal-epidermal junction. Bar graphs show mean ± SD. Individual dots represent average measurements from 4-6 HPFs per wound from different animals. \**P* < .05, \*\**P* < .01, \*\*\**P* <.001.

### Magnesium Treatment Improved Neovascularization

We tested the effect of magnesium wire treatment on capillary lumen density (neovascularization) in the wound bed at day 7 by staining wound cross sections with antibodies against CD31, or platelet endothelial cell adhesion molecule 1 (PECAM-1), a marker for endothelial cells. Imaging of the cross sections showed increased capillary lumen density in the granulation tissue of magnesium wounds as compared to PBS treatment **(Figure 3a-b).**

**Figure 3.**
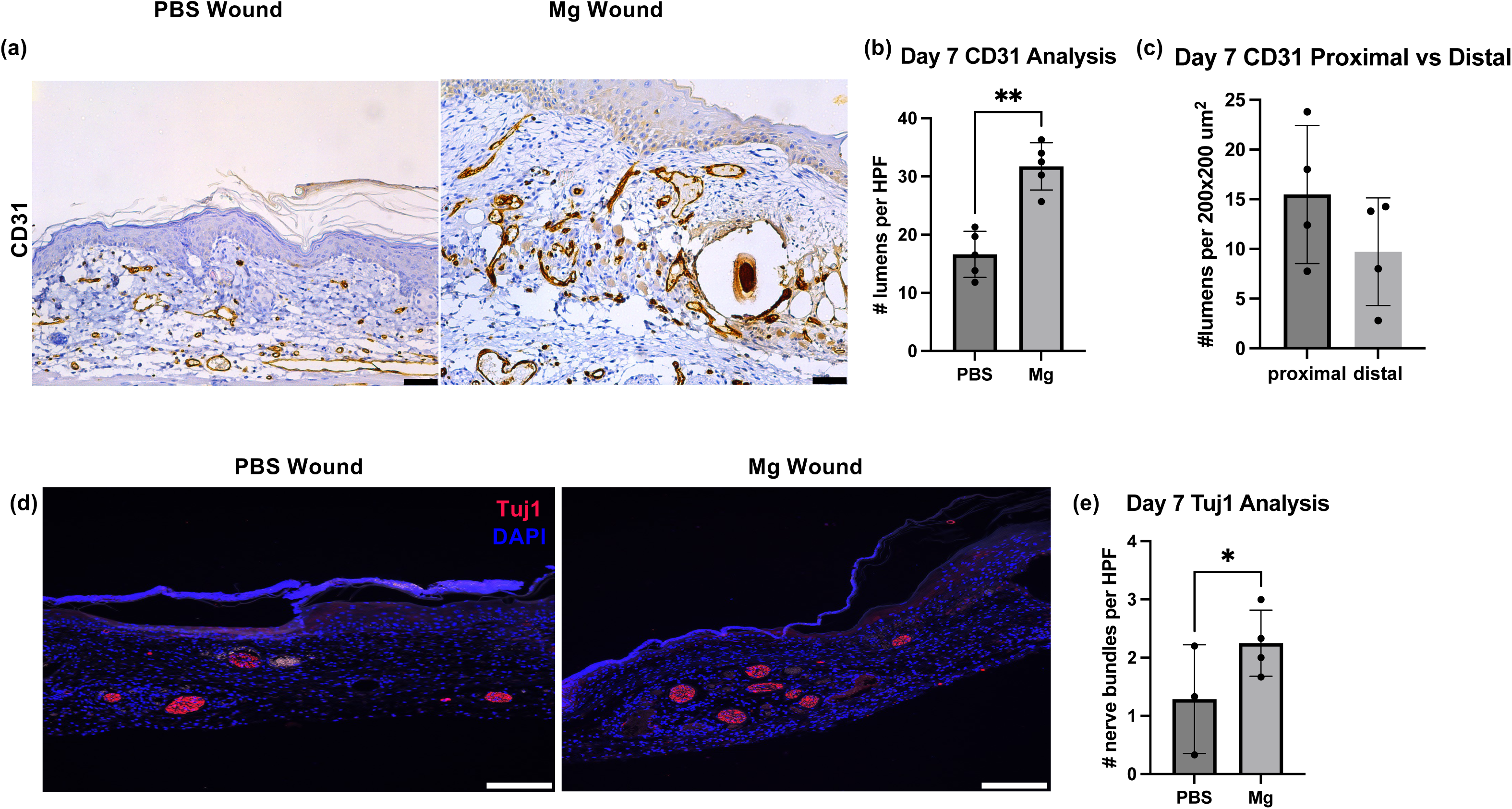
Mg increases capillary lumen density. (a) CD31 (endothelial cells) IHC staining of PBS and Mg wounds at day 7 post-wounding was performed and 20x representative images of wounds are shown. Scale bar = 50 µm. (b) Quantification of lumens per HPF from 4-6 HPFs across the wound bed per wound (40x). (c) Quantification of lumens proximal (within 200 µm from Mg wires) vs distal (200 µm away from Mg wires) at day 7 post-wounding. (d) Tuj1 stained positive nerve bundles in PBS and Mg wounds at day 7 post-wounding. Scale bar = 200 µm. (e) Quantification of % of Tuj1 positive nerve bundles per HPF (40x) at day 7 post-wounding. Bar graphs show mean ± SD. Individual dots represent average measurements from 4-6 HPFs per wound from different animals. \**P* < .05, \*\**P* < .01, \*\*\**P* < .001.

Analysis demonstrated significantly increased CD31-positive lumens in magnesium wire-treated wounds compared to PBS wounds (Mg 31.75 ± 4.06 vs PBS 16.62 ± 3.96 lumens per HPF; mean difference 15.13 lumens per HPF; 95% CI 6.72 – 23.54; *P* < .01). We further analyzed differences in lumen density in the wound bed depending on proximity to the magnesium wires by counted the number of lumens in 200 μm x 200 μm fields abutting the magnesium wires (proximal) and comparing it to the number of lumens in fields of the same size but at a further distance from the wire (distal) **(Supplemental Figure S1c)**. Analysis showed more vessels in the areas immediately surrounding the magnesium wires compared to more distal areas, however the difference was not statistically significant (proximal 15.49 ± 6.95 vs distal 9.71 ± 5.42 lumens per area; mean difference -5.77 lumens per area; 95% CI -18.80 – 7.25; *P* = .25, d = 0.71) **(Figure 3c)**.

### Magnesium Treatment Improved Nerve Repair

To determine the effect of magnesium on nerve repair, the cross sections of PBS- and magnesium wire-treated wounds at day 7 post-wounding were stained with Tuj1, an antibody against class III beta-tubulin, which was used as a neuronal marker. Slides appeared to have more nerve bundles and increased speckles of Tuj1-positive staining in the magnesium-treated wound bed **(Figure 3d-e).** Upon quantification, there were 74% more Tuj1 bundles in magnesium wire-treated wounds (Mg 2.25 ± 0.57 vs PBS 1.29 ± 0.93 Tuj1+ nerve fibers per HPF; mean difference 1.05 fibers per HPF; 95% CI 0.37 – 1.72; *P* = .02).

### Magnesium Treatment Reduced Scar Formation and Improved Tissue Regeneration

We followed wound healing until 28 days post-wounding. Gross imaging of the completely healed skin wounds at day 28 post-wounding showed minimal scarring in the magnesium wire-treated wounds compared to PBS controls **(Figure 4a).** Micro-CT images of the wounds taken at day 28 showed wire degradation within the wounds, which supports their safety and bioabsorption for use in wound healing applications **(Figure 4b)**. Trichrome staining of the wound sections was performed to evaluate the effect of treatments on ECM collagen remodeling, which showed better ECM organization and a basketweave pattern of collagen in magnesium wire-treated wounds compared to PBS controls **(Figure 4c).** Scar area at day 28 was measured using H&E cross sections. The scar area appeared smaller in magnesium wire-treated wounds; however, the difference was not statistically significant (Mg 0.28 ± 0.21 vs PBS 0.58 ± 0.20 mm^2^; mean difference -0.30 ± 0.17 mm^2^; 95% CI -0.77 – 0.16; *P* = .14, d = 0.29) **(Figure 4d).** Collagen content was further assessed from picrosirius-stained scar images. Although there appeared to be more collagen in magnesium scars than PBS scars, the difference was not significant (Mg 40.98 ± 12.09 vs PBS 23.68 ± 0.85; mean difference -30 ± 0.17; 95% CI -0.77 – 0.16; *P* = .07, d = 2.02) **(Figure 4e-f)**.

**Figure 4.**
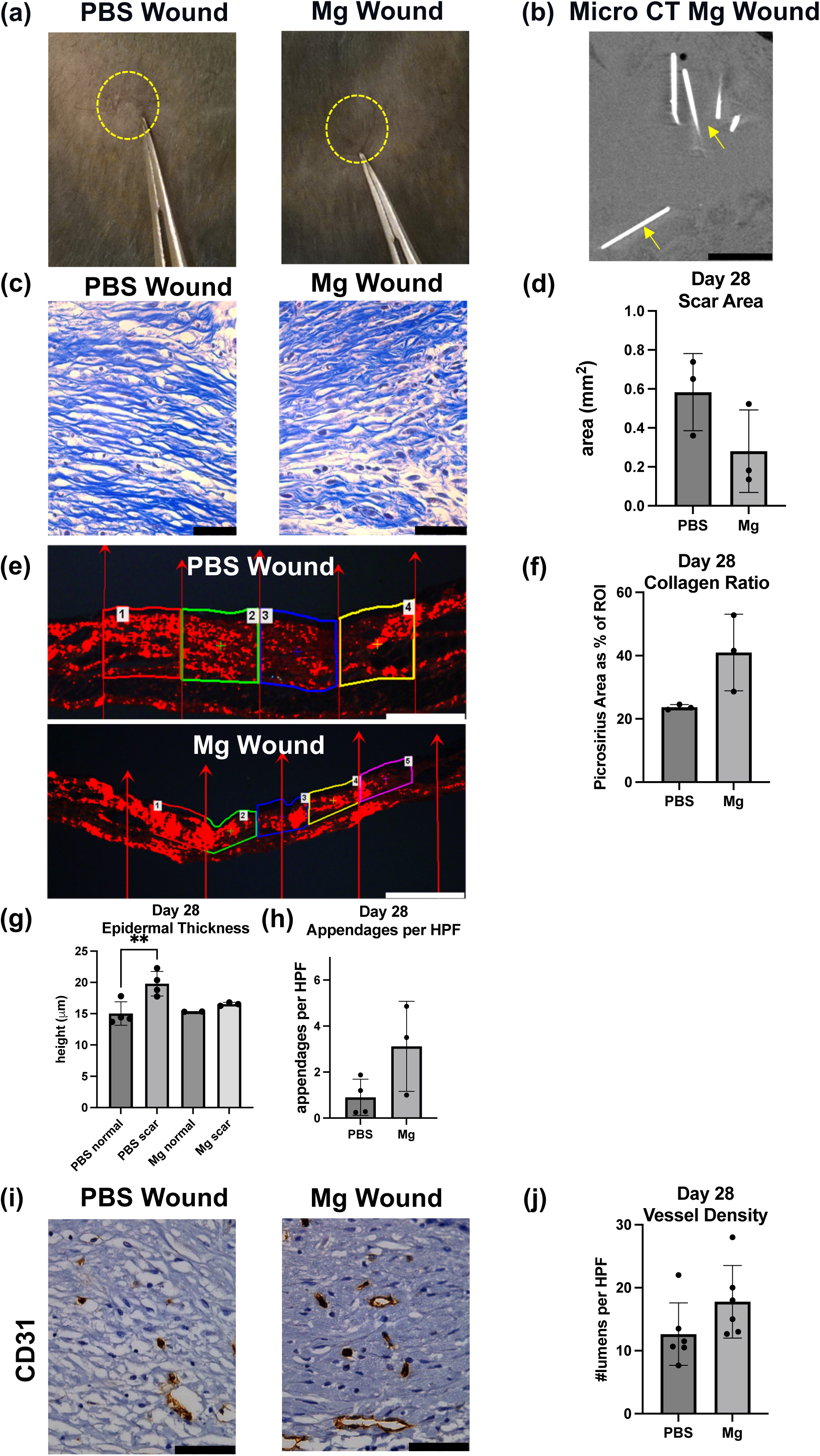
Mg improves tissue regeneration. (a) Pictures of scars of PBS and Mg wounds at day 28 post-wounding. Yellow dashed lines indicate scar area. (b) Mg wires (yellow arrows) in skin explants on micro-CT at day 28. Scale bar = 2.5 mm. (c) Trichrome staining of Mg wound shows basket weave pattern compared to parallel collagen bundles in PBS wound. Scale bar = 50 µm. (d) Scar area measured at day 28. (e) Picrosirius red fluorescence in PBS and Mg wounds at day 28. Scale bar = 100 µm. (f) The scars were divided into 4 parts to represent the middle area and edges, and picrosirius positive area was calculated as percent of region of interest (ROI) area to represent collagen ratio in PBS and Mg wounds at day 28. (g) Epidermal thickness at day 28. (h) Dermal appendages/HPF at day 28. (i) CD31 IHC staining of Mg and PBS wounds at day 28, Scale bar = 50 µm. (j) Vessel density at day 28 determined by CD31 staining. Bar graphs show mean ± SD. Individual dots represent average measurements from 4-6 HPFs per wound from different animals. \**P* < .05, \*\**P* < .01, \*\*\**P* < .001.

Further morphometric analyses of tissues taken at 28 days post-wounding were undertaken to quantify epidermal thickness, dermal appendages per HPF, and CD31 staining for blood vessel lumen density using H&E sections. The PBS scars had a thicker epidermis in the scar area compared to adjacent normal skin (PBS scar 19.8 ± 1.97 μm vs PBS normal 15.03 ± 1.88 μm; mean difference 4.77; 95% CI 2.55 – 6.99; *P* < .01), whereas the scars in magnesium wire-treated wounds had similar thickness of the epidermis as the normal skin (Mg scar 16.53 ± 0.28 μm vs Mg normal 15.33 ± 0.02 μm; mean difference 1.21; 95% CI -2.16 – 4.57; *P* = .14) **(Figure 4g)**. This indicates efficient tissue repair with magnesium, but not PBS, as evidenced by epidermal thickness returning to normal in the scar. While there were more dermal appendages (Mg 3.12 ± 1.96 vs PBS 0.90 ± 0.78 appendages per HPF; mean difference 2.22 ± 1.05 appendages per HPF; 95% CI -0.49 – 4.92; *P* = .09, d = 1.61) **(Figure 4h)**, and more blood vessels (Mg 17.78 ± 5.76 vs PBS 12.64 ± 4.96 lumens per HPF; mean difference 6.97 lumens per HPF; 95% CI -0.70 – 14.63; *P* = .07, d = 1.35) **(Figure 4i-j)** in magnesium wire-treated wounds, these measures were not significantly different. There also were increased subepithelial nuclei in magnesium scars, although the increase did not reach significance (Mg scar 6559.22 ± 1150.43 vs PBS scar 4715.75 ± 630.46 nuclei per mm^2^; mean difference 1843 ± 757.4 nuclei per mm^2^; 95% CI -259.4 – 3946; *P* = .07, d = 2.0), and there were differences between the scar and normal tissues for both magnesium (Mg scar 6559.22 ± 1150.43 vs Mg normal 2105.39 ± 234.47 nuclei per mm^2^; mean difference 4454 nuclei per mm^2^; 95% CI 1527 – 7381; *P* = .02) and PBS (PBS scar 4715.75 ± 630.46 vs PBS normal 2161.67 ± 834.66 nuclei per mm^2^; mean difference 2554; 95% CI -373.5 – 5482; *P* = .06, d = 2.17). This suggests an as-yet uncharacterized difference in cellularity, presumably changes in fibroblasts **(Supplemental Figure S2)**.

### Degrading Magnesium Wires Influence Wound Healing Mechanics

The complex *in vivo* degradation process of the magnesium wires influences wound healing mechanics by introducing strain gradients in the tissue during movement or postural changes **(Figure 5a)**. Two representative deformation modes were modeled using finite element analysis (FEA) to quantify strain in the wires over the course of healing, expressed as percentage of wound closure **(Figure 5b)**. The chemo-mechanical analysis was conducted in two parts: a chemical analysis to track biofluid concentration in the wires and a mechanical analysis to evaluate changes in Young’s modulus induced by the corrosive conditions. Wire degradation is initially approximated using a first-order kinetics model to estimate an upper bound on the rate of magnesium degradation, which in the physiological environment is highly sensitive to water and salt concentrations, oxygen levels, pH, and temperature. This approach also captures the resulting nonlinear concentration profile within the wires, followed by transient mechanical effects causing modulus reduction **(Figure 5c)**. Under uniaxial stretching, strain in the degrading wires increased monotonically, reaching 0.2%, while non-degrading wires remained near 0.05%. In bending, early tissue remodeling (< 25% wound closure) initially reduced wire strain, followed by an increase to 0.68% in degrading wires compared to 0.13% in non-degrading wires. **Figure 5d** shows simulations of tissue deformation and strain evolution in the wound, surrounding tissue, and degrading wires. Under 10% uniaxial stretching along the wire axis, strains up to 50% developed at the wire-wound interfaces. Parametric studies of magnesium wire radius and spacing **(Supplemental Figure S3)** show that increasing the wire radius from 107 µm to 167 µm reduces maximum strain by 23% under bending and 40% under stretching, whereas changes in inter-wire spacing have a comparatively minor effect (5% change).

**Figure 5.**
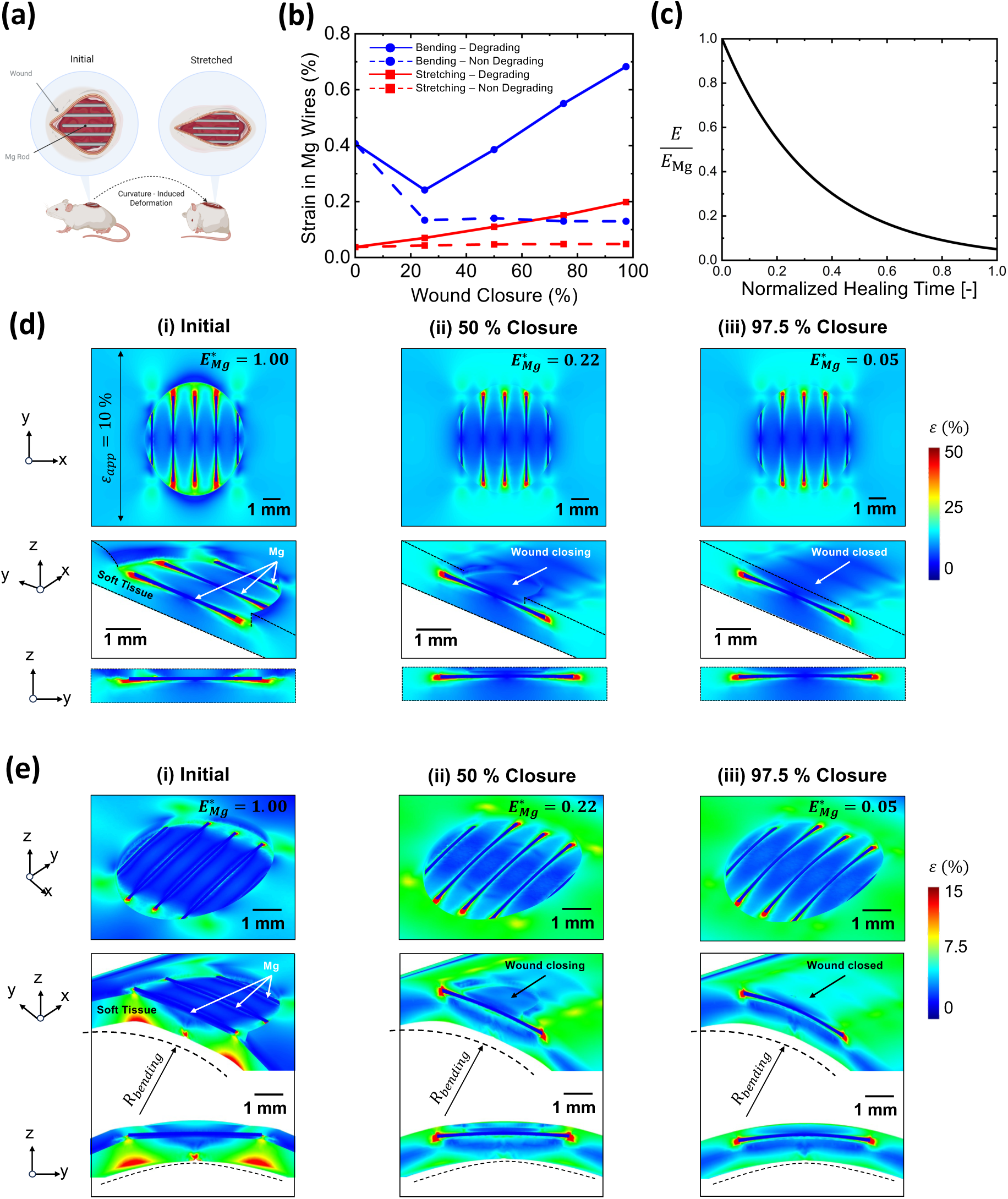
Mg introduces strain gradients under deformation. (a) Illustration of the wound with implanted magnesium wires showing curvature-induced deformations from mouse posture. Created with Biorender. (b) Maximum strain in magnesium wires after stretching and bending for degrading and non-degrading material models. (c) Modulus ratio of degrading magnesium wires as a function of the wound healing time. Simulation results showing strain distributions in the wound under (d) uniaxial deformation and (e) bending during progressive wound closure (i–iii, 0–97.5%). For tissue material properties, data from 12 week dorsal skin of female mice was utilized.^41–42^

As the wound closed **(Figure 5e, panels i–iii)**, strain around the wires and at the new wound surfaces redistributed, reducing skin surface strains to below 10%. **Figure 5e** shows strain in the wound under a bending radius of 13.5 mm, with maximum strains reaching 15% near the magnesium wires. Tissue remodeling, modeled as isotropic wound closure, significantly altered strain redistribution during bending at 50% and 97.5% closure, affecting not only the surface but also the tissue thickness. Combined tissue structural changes and magnesium degradation governed the formation of strain gradients, with deformation-dependent patterns that may locally affect mechanical signaling during healing. **Figure 6** shows the normalized change in Young’s modulus *E/E*_Mg_, representing the progressive non-linear mechanical loss of magnesium within the wound environment. The ratio starts at *E/E*_Mg_ =1 at the onset of healing and exponentially declines to approximately 0.05 as the wound fully closes, reflecting substantial structural loss of the magnesium wire.

**Figure 6.**
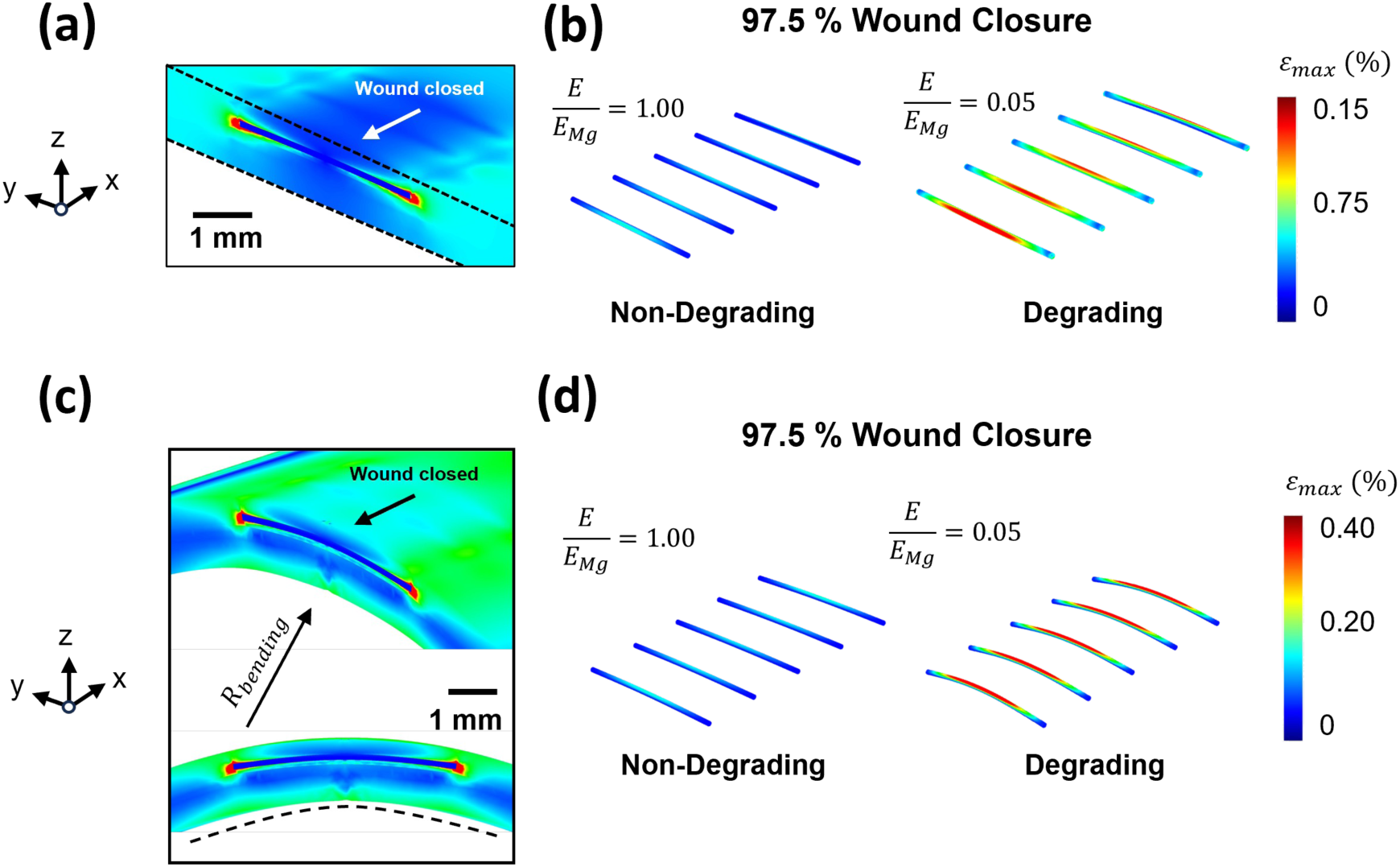
Strain distributions in wound tissues and magnesium wires under (a, b) uniaxial stretching and (c, d) bending. For each loading condition, panels show (a, c) strain in the wound tissues and (b, d) strain in the five magnesium wires for non-degrading and degrading cases at near-complete wound healing.

## DISCUSSION

This study demonstrates that our previously characterized bioabsorbable magnesium metal wires enhance wound healing in full-thickness murine dermal wounds through immunoregulatory, angiogenic, and mechanical effects. Magnesium wire-treated wounds exhibited increased granulation tissue, greater vascular density, and more nerve bundles, aligning with prior studies suggesting magnesium supports regenerative healing.^10^ Importantly, the wires did not trigger a foreign body response, which can be detrimental to healing. Minimal collagenous encapsulation of the wires and reduced infiltration of F4/80-postive macrophages and CD45-positive leukocytes suggest an anti-inflammatory effect of magnesium.

Magnesium ion levels peak during the proliferative phase of wound healing in rodents, potentially supporting the increased cell metabolism and proliferation needed for repair.^2^ Consistent with this, magnesium-treated wounds displayed more granulation tissue at day 7.^9^ Neovascularization was significantly enhanced, likely supporting improved granulation and tissue repair.^9,41,42^ While more vessels appeared near the wires (proximal), the difference was not statistically significant, potentially due to coverage of the wound area by the wires and even ion distribution. Neuronal staining and nerve bundle density were also increased in magnesium-treated wounds, suggesting possible neurotrophic effects that warrant further study.

By day 28, magnesium-treated wounds had smaller scars and improved collagen architecture, with trichrome staining revealing a more organized, basketweave collagen pattern. While neither the reduction in scar size (*d* = 0.29) nor the increase in collagen content (*d* = 2.02) reached statistical significance, the effect sizes provide important context. The small effect size for scar area suggests only a modest reduction, likely limited by biological variability and small sample size. In contrast, the very large effect size for collagen content indicates a substantial magnitude of difference despite the *P* value not crossing the significance threshold, consistent with qualitative improvements in ECM organization. Together, these trends support the possibility that magnesium enhances late-stage matrix remodeling even when quantitative comparisons are underpowered. Micro-CT confirmed wire degradation, with small fragments remaining in the wound, suggesting ongoing ion release without adverse effects. While all morphometric analyses showed visible improvement with magnesium treatment, only the reduction in epidermal thickness, signifying faster restoration of normal epidermal structure, reached statistical significance. Further studies should evaluate restoration of barrier function (e.g., transepidermal water loss), as defective barrier recovery marks impaired wound healing.

Computational modeling showed that wire degradation dynamically increased strain gradients during healing. Compared to inert wires, degrading wires imposed greater mechanical loading on the tissue over time, particularly under stretch and bending. These mechanical cues may influence fibroblast behavior, ECM remodeling, and scar architecture. This dual biochemical and biomechanical modulation highlights magnesium’s potential as a regenerative scaffold component. These findings underscore the importance of implant degradation kinetics and tissue mechanical adaptation in wound repair. Magnesium wires provide a promising platform for regenerative dermal healing and may be applicable to broader surgical contexts, owing to their ease of manufacturing into lattices and sheets with tunable porosity and thickness. Future studies integrating molecular readouts with biomechanical modeling are warranted to better understand the role of dynamic strain gradients in guiding regenerative healing in wire-treated wounds.

Our results complement prior work demonstrating that magnesium promotes wound closure, neovascularization, and collagen production. However, earlier studies using polymer-based or composite magnesium systems introduced confounding materials and degraded rapidly *in vivo*.^8,15–20^ Other recent studies have explored magnesium-based microneedle systems and hybrid scaffolds to support wound healing. These platforms, which often use dissolvable polymers or composites, allow for controlled, localized delivery of magnesium ions and have demonstrated accelerated wound closure, enhanced angiogenesis, and immunomodulation in various models.^43^ For example, magnesium-loaded microneedle patches have been shown to promote angiogenesis and facilitate tissue regeneration, similar to the effects observed in our model.^8,44^ However, potential confounding due to polymer degradation and rapid magnesium release remains a limitation of these systems. Additionally, while microneedle systems can conveniently deliver magnesium to the dermis, our wires have the potential to not only be used in the epidermis, but to be spun into different scaffolds to use across a wide range of applications. Our approach, utilizing pure bioabsorbable magnesium wires, also differs in providing sustained ion release due to slower degradation and adding mechanical support while minimizing the introduction of exogenous polymers, thus offering an alternative strategy for prolonged and biometric regenerative support.

Our study has several limitations. Testing was done in a murine model, and while well-suited for mechanistic studies, murine skin has important differences with human skin. Additionally, the study follow-up was limited to 28 days, and long-term outcomes such as complete tissue remodeling and restoration of normal skin architecture were not assessed. , While we did see a decrease in inflammatory cells in magnesium-treated wounds, prior literature has reported that magnesium ions promote shifts toward reparative macrophage phenotypes.^45^ We did not assess macrophage polarization, and future studies should include immunophenotyping to validate this mechanism. Functional outcomes, such as tensile strength, barrier recovery, and mechanical integrity of the healed tissue were not measured. Finally, although micro-CT imaging and finite element modeling provided insight into degradation and strain dynamics, these models may not fully capture *in vivo* corrosion kinetics or the complex host-material interaction occurring over time. Direct *in situ* strain measurements for the implanted magnesium wires were not available; therefore, the finite element model was verified via mesh convergence and internal consistency mechanics checks, and was constructed to mirror the experimental implantation geometry, materials, and expected loading conditions. Magnesium ions, crucial for cellular functioning, can reach toxic levels if applied to cells in culture, however metal implants in rodent and humans have almost no magnesium-derived detrimental effects and the metal degrades safely.^33^ However, a limitation is that selective magnesium ion detection is not possible *in vivo* so novel methods are required to detect magnesium implants.^46^ Future studies incorporating molecular profiling, mechanical testing, and large-animal models will be critical to validate our findings, describe tissue compatibility, and guide clinical translation.

Overall, bioabsorbable magnesium wires enhanced wound closure, neovascularization, granulation tissue formation, and reduced inflammation at early time points in full-thickness murine wounds, while promoting improved ECM remodeling and normalization of scar epidermal thickness at later stages. These results support continued development of magnesium-based scaffolds for surgical wound care and regenerative medicine applications.

## KEY FINDINGS

- In a murine full-thickness dermal wound model, bioabsorbable magnesium wire-treated wounds showed significantly improved granulation tissue formation, reduced inflammation, and increased neovascularization by day 7
- By day 28, magnesium-treated wounds showed improved collagen organization and normalized epidermal thickness
- Magnesium wires degraded *in vivo* without eliciting a foreign body response and computational modeling revealed dynamic strain gradients introduced during degradation that may further influence regenerative signaling

## ACKNOWLEDGMENTS

This project was supported by biostatistician Keely Wolf of the Baylor College of Medicine Michael E. DeBakey Department of Surgery. We also acknowledge support for this research by the NIH/NIGMS R01GM141366 (SB), John S. Dunn Foundation Collaborative Research Award Program administered by the Gulf Coast Consortia (SB), Clayton Seed Grant by the Department of Pediatric Surgery at Texas Children’s Hospital (SB), and RSI awards seed funding from Rice (SB and RA).

Declaration of Competing Interest: The authors declare that they have no competing interests.

Meeting Presentation: This work was presented at the 19th Annual Academic Surgical Congress in February 2024, in Washington D.C. and 2024 WHS (Wound Healing Society) Meeting in May 2024 in Florida.

## AUTHOR CONFIRMATION

We, the authors, confirm that this manuscript entitled “Bioabsorbable Magnesium Metal Scaffolds Improve Dermal Wound Healing and Tissue Regeneration” is original, has not been published elsewhere, and is not under consideration by any other journal. All authors have contributed substantially to the conception, design, data acquisition, or analysis and interpretation of data, have been involved in drafting or revising the manuscript, and have approved the final version for submission. All authors agree to be accountable for all aspects of the work and affirm that any questions related to the accuracy or integrity of any part of the work will be appropriately investigated and resolved. All listed authors meet the criteria for authorship, and no individuals who do not meet these criteria have been included as authors. All animal procedures were conducted in accordance with institutional and national ethical standards and were approved by the Cincinnati Children’s Hospital Medical Center Institutional Animal Care and Use Committee (IACUC). This manuscript is being submitted solely to *Advances in Wound Care*.

## Contributions

SB and SP designed the study; SB, NNA, SP, TH, XA designed and performed experiments; MEG, NNA, AN, LY, SK, XA, SP analyzed data; PL, CL, RA performed mechanical modeling; MEG, NNA, RA, made the figures; MEG, NNA, RA, SB, SP drafted/revised the paper; SB and SP provided scientific comments for the paper; all authors discussed the results and commented on the manuscript.

## AUTHOR DISCLOSURE AND GHOSTWRITING

The authors declare that no ghostwriting or undisclosed writing assistance was used in the preparation of this manuscript. All authors contributed directly to the writing and critical revision of the article.

## FUNDING STATEMENT

This work was supported by Grant Number R01GM141366-02A from NIH/NIGMS (SB), a grant from the National Sciences Foundation Engineering Research Center for Revolutionizing Metallic Biomaterials (EEC-0812348, NCAT 260116C supplement) (SP, TH, XA), the John S. Dunn Foundation Collaborative Research Award Program administered by the Gulf Coast Consortia (SB), a Clayton Seed Grant by the Department of Pediatric Surgery at Texas Children’s Hospital (SB), and RSI awards seed funding from Rice University (SB and RA).

## ABOUT THE AUTHORS

Mary Elizabeth Guerra is a surgery resident at Baylor College of Medicine (BCM) and a research fellow in the Laboratory for Regenerative Tissue Repair (LRTR). (first author) Nabila Anika is a surgery resident at the USF Health Morsani College of Medicine.

Ajeet Nagi is an undergraduate student at The University of Texas at Austin.

Tracy Hopkins is a Senior Research Assistant in Neurosurgery at the University of Cincinnati. Xiaoxian An is a Research Associate at Weill Cornell.

Ling Yu is a Research Assistant in the LRTR.

Pei Liu and Chihtong Lee are PhD students at Rice University.

Sundeep Keswani is a Professor of Surgery at BCM and LRTR co-director, whose research explores inflammation-ECM interactions in wound healing.

Raudel Avila, Assistant Professor of Mechanical Engineering at Rice University, directs the Computational Mechanics and Bioelectromagnetics Laboratory, developing models of soft and transient electronic systems.

Sarah Pixley is an Associate Professor in the Department of Pharmacology, Physiology, and Neurobiology at the University of Cincinnati, studying tissue reactions biodegradable metal implants.

Swathi Balaji is an Associate Professor at BCM and LRTR director. She has expertise in mechanisms behind dermal fibrosis and wound healing. (last author)

## ABBREVIATIONS AND ACRONYMS

d7: day 7
d28: day 28
ECM: extracellular matrix
ED: epidermis-dermis
FBR: foreign body response
FEA: finite element analysis
H&E: Hematoxylin and Eosin
IHC: immunohistochemistry
Mg: magnesium
Micro-CT: microcomputed X-ray tomography
PBS: phosphate-buffered saline
PC: Panniculus carnosus
ST: subcutaneous tissue
WT: wildtype

---

**Supplemental Figure S1.** (a) Methods and study timeline. Gross image of stented wound with magnesium wires in place. (b) Wound morphometry and IHC analysis. Solid vertical arrows represent original wound margins and dotted lines represent epithelial margins (extent of re-epithelialization). Thin black boxes represent evenly spaced 40x HPFs. (c) Assessment of vessel density proximal (surrounding) and distal (adjacent) to Mg wires at day 7. Asterisks (*) indicate locations of Mg wires. Scale bar = 250 µm.

**Supplemental Figure S2.** Increased subepithelial nuclei in Mg wounds. (a) Fluorescent imaging of day 28 wounds. Nuclei stained with DAPI. Scale bar = 30 µm. (b) Quantification of #subepithelial nuclei per mm^2^. \**P* < .05, \*\**P* < .01, \*\*\**P* < .001.

**Supplemental Figure S3.** (a) Finite element mesh-convergence analysis demonstrating stabilization of strain-energy predictions across coarse and refined meshes. (b) Maximum strain in Mg wires as a function of wire radius under both stretching and bending deformations. (c) Maximum strain in Mg wires as a function of wire spacing under stretching and bending, highlighting geometric sensitivities in mechanical loading.

## Notes

### Competing Interest Statement

The authors have declared no competing interest.

