## Supplementary figures and images for "Bioabsorbable Magnesium Metal Scaffolds Improve Dermal Wound Healing and Tissue Regeneration"

### Supplemental Figure S1

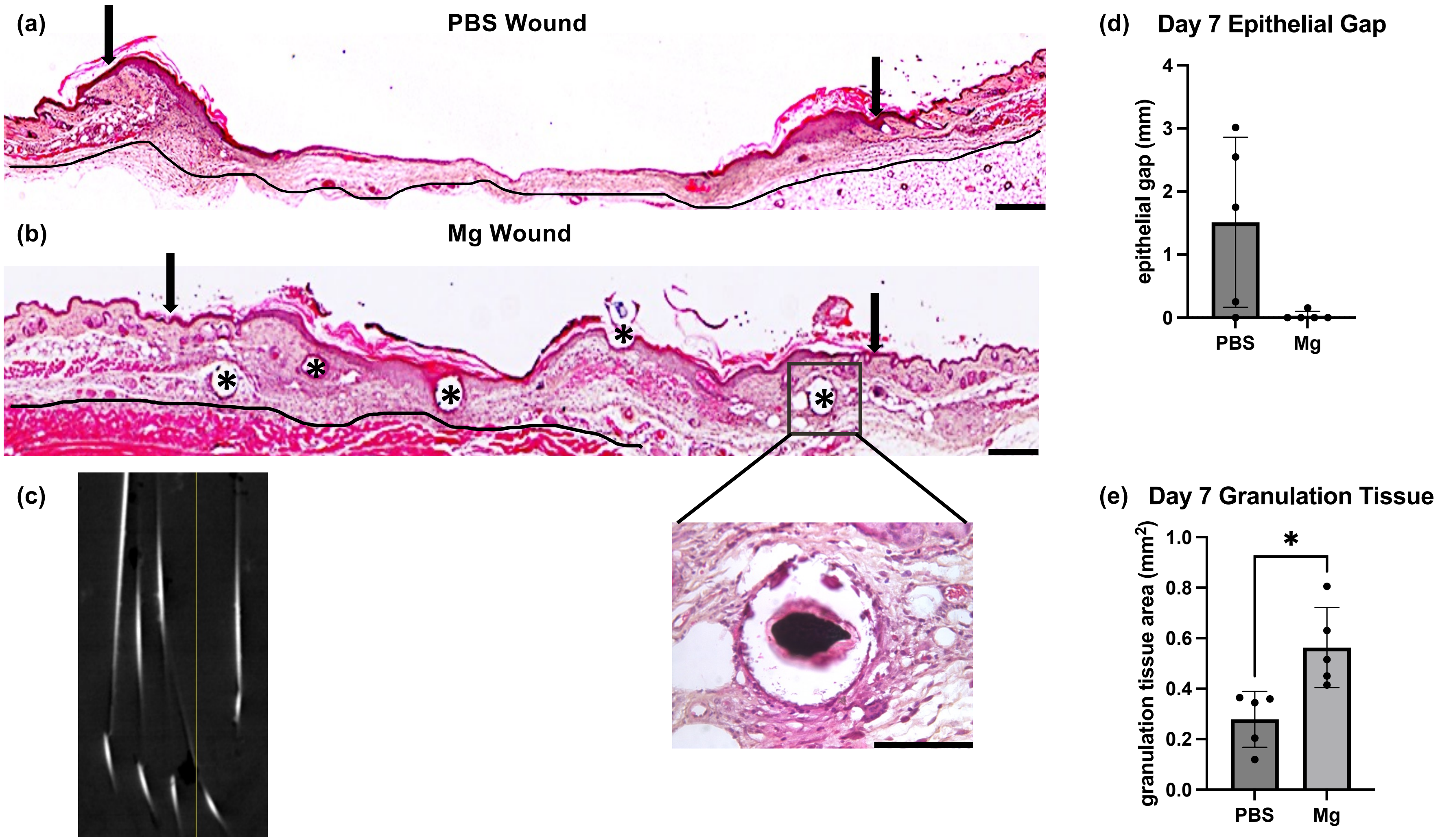

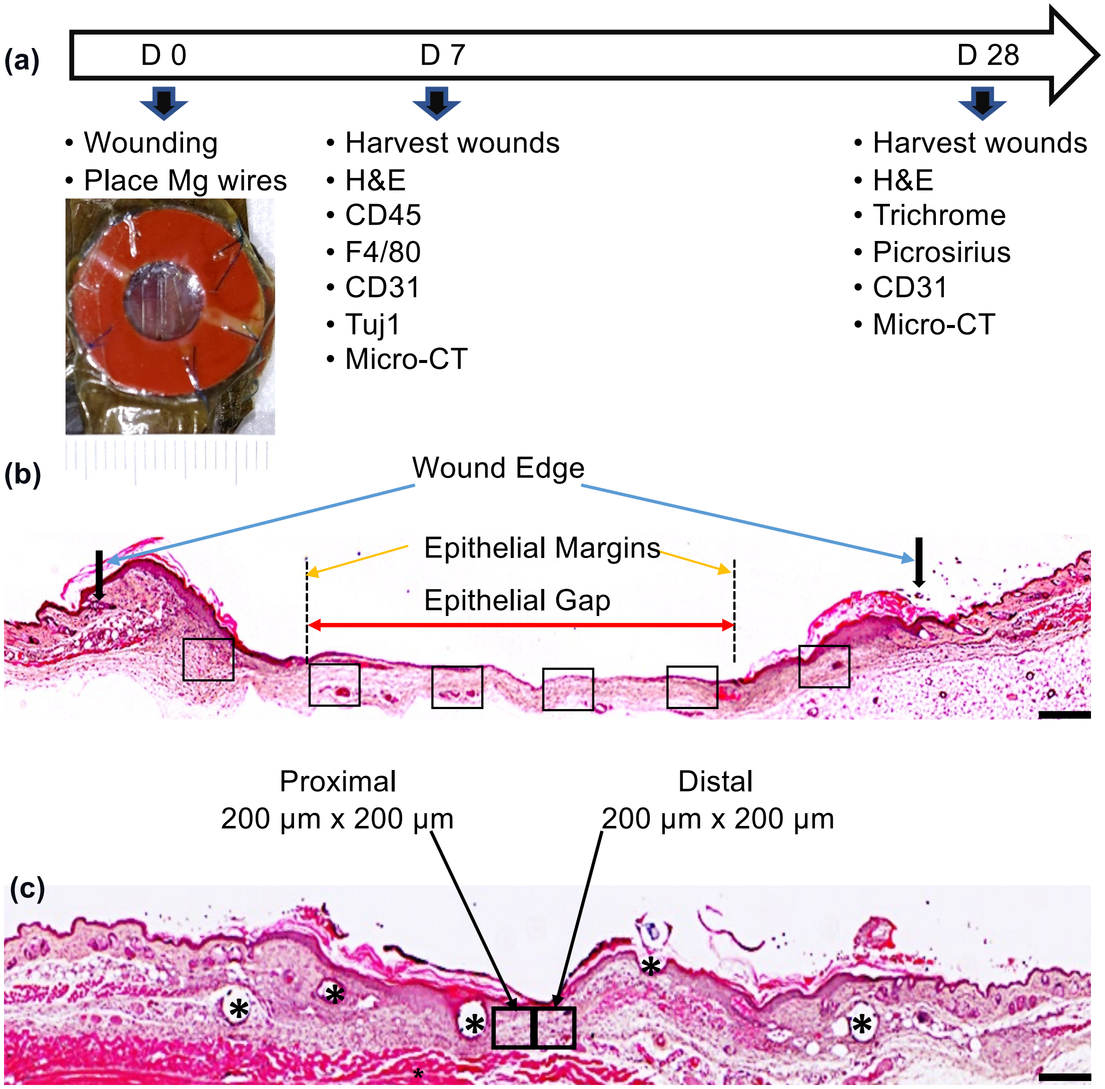

### Supplemental Figure S2

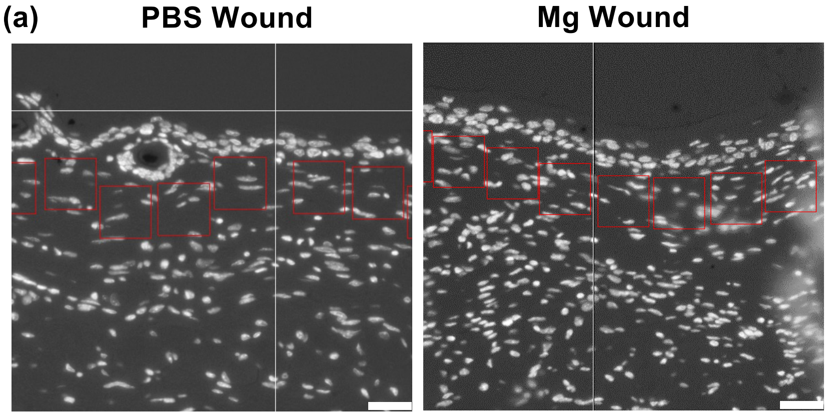

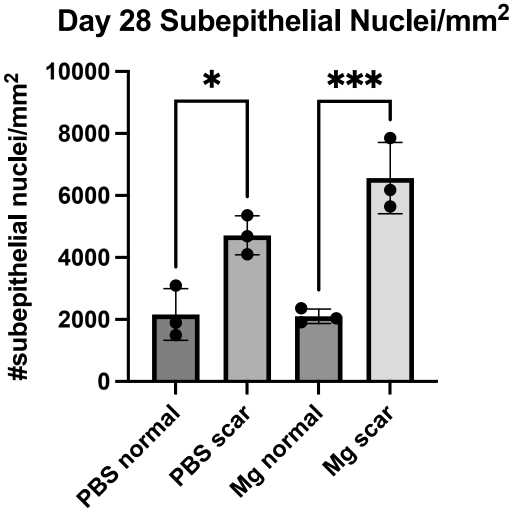

### Supplemental Figure S3

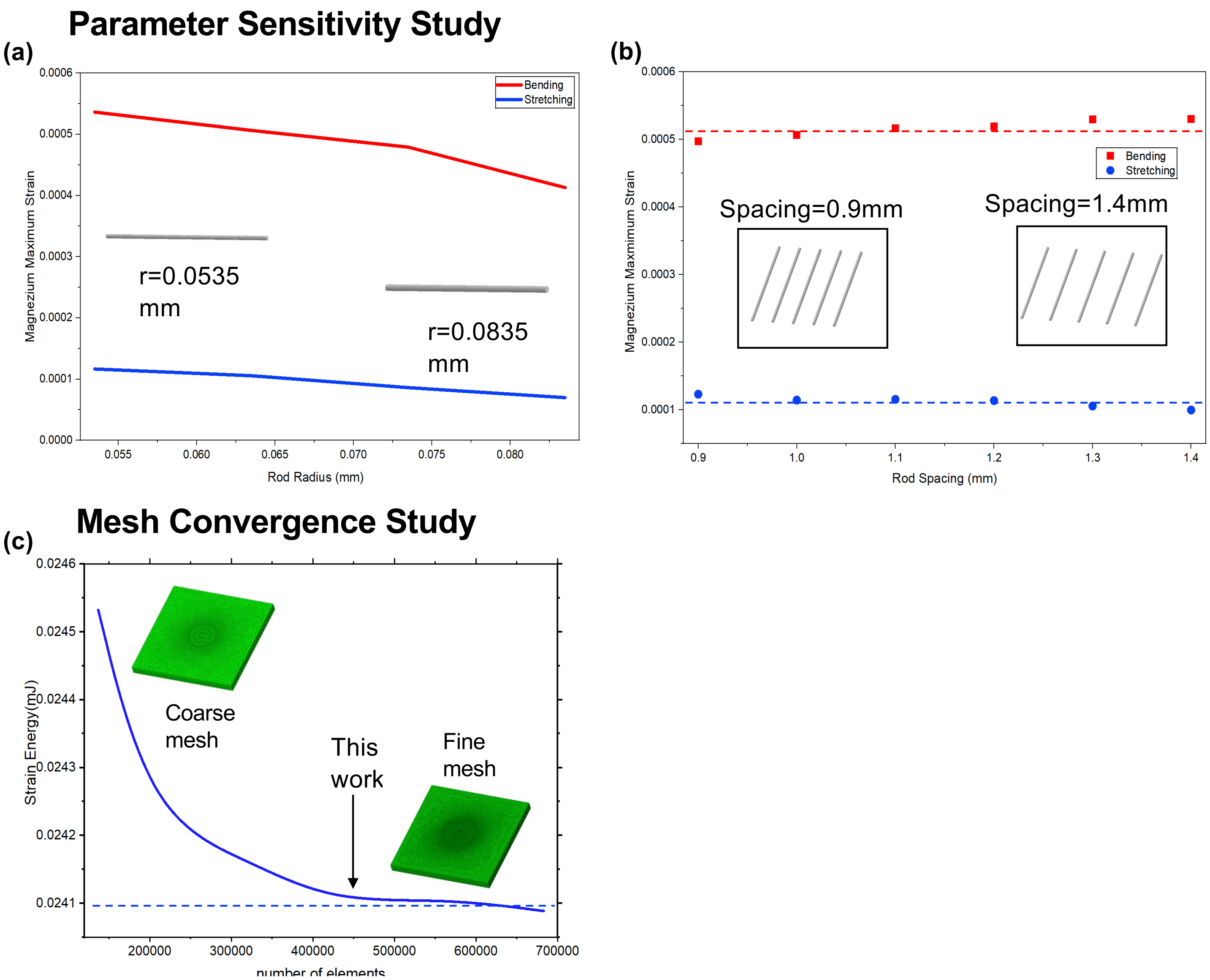
